# Conserved molecular signatures of hygrosensory neurons in two dipteran species

**DOI:** 10.1101/2025.03.17.643622

**Authors:** Kristina Corthals, Ganesh Giri, Johan Reimegård, Allison Churcher, Anders Enjin

## Abstract

Small poikilothermic animals like insects rely on environmental sensing for survival. The ability to detect humidity and temperature through specialized sensory neurons is particularly critical, allowing them to maintain water balance across diverse environments. While recent studies have identified key receptors associated with humidity sensing, our understanding of the broader molecular architecture underlying these sensory systems remains incomplete. Here, we conducted a comparative analysis of humidity receptor neurons (HRN) between the vinegar fly *Drosophila melanogaster* and the yellow fever mosquito *Aedes aegypti*. We identified 21 genes that define the molecular identity of HRNs. These genes encode proteins involved in transcriptional regulation, cellular signaling, enzymatic pathways and cellular organization. Through behavioral analysis, we demonstrate essential roles for three of these genes, the serotonin receptor 5-HT7, the transcription factor nubbin and the kinesin motor protein Kif19A are all required for humidity-guided behavior. The conservation of this molecular toolkit between species separated by over 200 million years of evolution suggests shared functional requirements for environmental sensing in insects. Our findings provide insights into fundamental principles of sensory neuron organization and offer a framework for understanding how specialized sensory systems evolve and maintain their function.

## Introduction

Environmental humidity fundamentally shapes terrestrial ecosystems and profoundly influences animal behavior and survival^1,2^. For insects, as small poikilothermic animals, the ability to detect and respond to environmental moisture levels is particularly crucial, affecting their distribution, survival, and ecological interactions across diverse habitats^3,4^. The capacity to sense humidity serves multiple vital functions across insect species. Pollinators such as the bumble bee *Bombus terrestris*, the white-lined sphinx moth *Hyles lineata* and the tobacco hawkmoth *Manduca sexta* use floral humidity gradients to locate and identify flowers^5–7^. In disease vectors like the yellow fever mosquito *Aedes aegypti* and the malaria mosquito *Anopheles gambiae*, humidity sensing plays a crucial role in both host-seeking and oviposition site selection ^8–10^. These mosquitoes utilize their hygrosensory systems to detect subtle moisture gradients surrounding human bodies, which serve as essential short-range cues for host location^8–10^.

Insects detect humidity through specialized sensory structures called hygrosensilla which are located on their antennae^11^. Each hygrosensillum houses a characteristic group of three humidity receptor neurons (HRNs), known as a hygrosensory triad. This triad consists of three functionally distinct neurons: a moist neuron that is activated when humidity increases, a dry neuron that responds when humidity decreases, and a hygrocool neuron that activates when temperature drops^12^. The organization of these hygrosensilla varies among insect species. In the *D. melanogaster*, multiple hygrosensilla are clustered together within an invagination on the posterior side of the antenna called the sacculus^13^. In contrast, in *Ae. aegypti*, individual hygrosensory triads are housed in separate invaginations, each connected to the antennal surface through a small pore^14^.

The molecular basis of humidity and temperature sensing in insects involves several members of the ionotropic receptor (IR) gene family. In *D. melanogaster*, Ir40a is expressed in dry-responsive neurons in both chamber I and II of the sacculus, while Ir68a is found in moist-responsive neurons^15–18^. Hygrocool cells show a chamber-specific expression pattern, with Ir21a expressed in chamber I and Ir40a in chamber II^19^. Additionally, Ir25a and Ir93a are expressed in all these sensory neurons. Behavioral studies using mutants have provided evidence for the roles of these where loss of either Ir25a or Ir93a disrupts both humidity and temperature sensing behaviors^15,17,20^. Ir40a and Ir68a mutants have impaired humidity sensing, while Ir21a mutant flies show normal humidity responses, they exhibit defective thermosensation, suggesting a specific role in temperature detection^15–18,21^.

This molecular mechanism appears to be evolutionarily conserved across mosquito species. Recent studies have shown that Ir93a is essential for both temperature and humidity sensing in mosquitoes, as Ir93a mutants fail to respond to either stimulus^10^. Similar to *D. melanogaster*, Ir21a mutant mosquitoes show specific defects in temperature sensing while maintaining normal humidity responses^22^, Ir40a is associated with dry air detection, Ir68a to humid air, and these neurons are necessary for both blood-feeding and oviposition behaviors^10,14^.

Across these species, the ionotropic receptors Ir93a, Ir40a, and Ir68a emerge as critical molecular components essential for functional hygrosensation. This conservation of ionotropic receptor-mediated humidity sensing suggests a fundamental evolutionary mechanism in dipteran insects. However, beyond these receptors, little is known about the broader molecular architecture that enables hygrosensory neurons to develop and function. In a previous study we identified distinct transcriptional profiles of hygrosensory neurons in the sacculus, revealing unique molecular signatures for dry, moist and hygrocool neurons, along with specialized support cells^23^. Here, we build upon these insights and present comparative transcriptomic analysis of hygrosensory neurons in *D. melanogaster* and *Ae. aegypti*, two species separated by over 200 million years of evolution, to identify a conserved molecular toolkit of 21 genes that define these specialized neurons. Among these, we demonstrate essential roles for three previously uncharacterized genes in humidity sensing: the serotonin receptor 5-HT7, the transcription factor nubbin, and the kinesin motor protein Kif19A. Our findings reveal fundamental organizational principles of hygrosensory neurons and provide insights into how specialized sensory systems are built and maintained across evolution.

## Methods and Materials

### Methods

#### Data availability

github.com/hygrosensation/ComparativeStudy

#### Transcriptome analysis

##### Data

To analyze the transcriptomic profiles of hygrosensory (HRN) and thermosensory (TRN) neuron populations in *D.melanogaster* antennae, we utilized the 10x Genomics antennal dataset from the recently published Fly Cell Atlas^24^. For Ae. aegypti, we used two recently published single-nucleus dataset of a Ae. aegypti antenna^25,26^. Given the potential discrepancies in gene annotations and processing methodologies between the two available Ae. aegypti datasets, we opted against their integration to preserve the integrity of our downstream analyses. This decision mitigates the risk of introducing biases or artifacts that could arise from merging datasets with unknown preprocessing differences. Consequently, we will treat and analyze these datasets independently, ensuring robust and reliable results for each set. Quality control and subsequent analyses were performed using the Seurat package Version 5.1.0^27^.

##### Extraction of neuronal cells

All datasets were previously dimensionally reduced and clustered. The *D. melanogaster* dataset was processed as described in^23^. Consistent with the classification of cell types in Corthals et al, 2023 clusters were assigned neuronal identity based on the expression of 4 neuronal marker genes.

Genes used were: Syt1, elav, CadN, brp (Supp Figure 1 A) and their *Ae. aegpyti* orthologues LOC5565901 (orthologue to Syt1), LOC5570204 (orthologue to elav), LOC5564848 (orthologue to CadN), LOC5570381 (orthologue to brp) (Supp Fig 2 A). Additionally, glia cells were classified using the *D. melanogaster* glial maker repo and the *Ae. aegypti* orthologue LOC110678282 (Supp Fig 1 B and 2 B).

**Figure 1.**
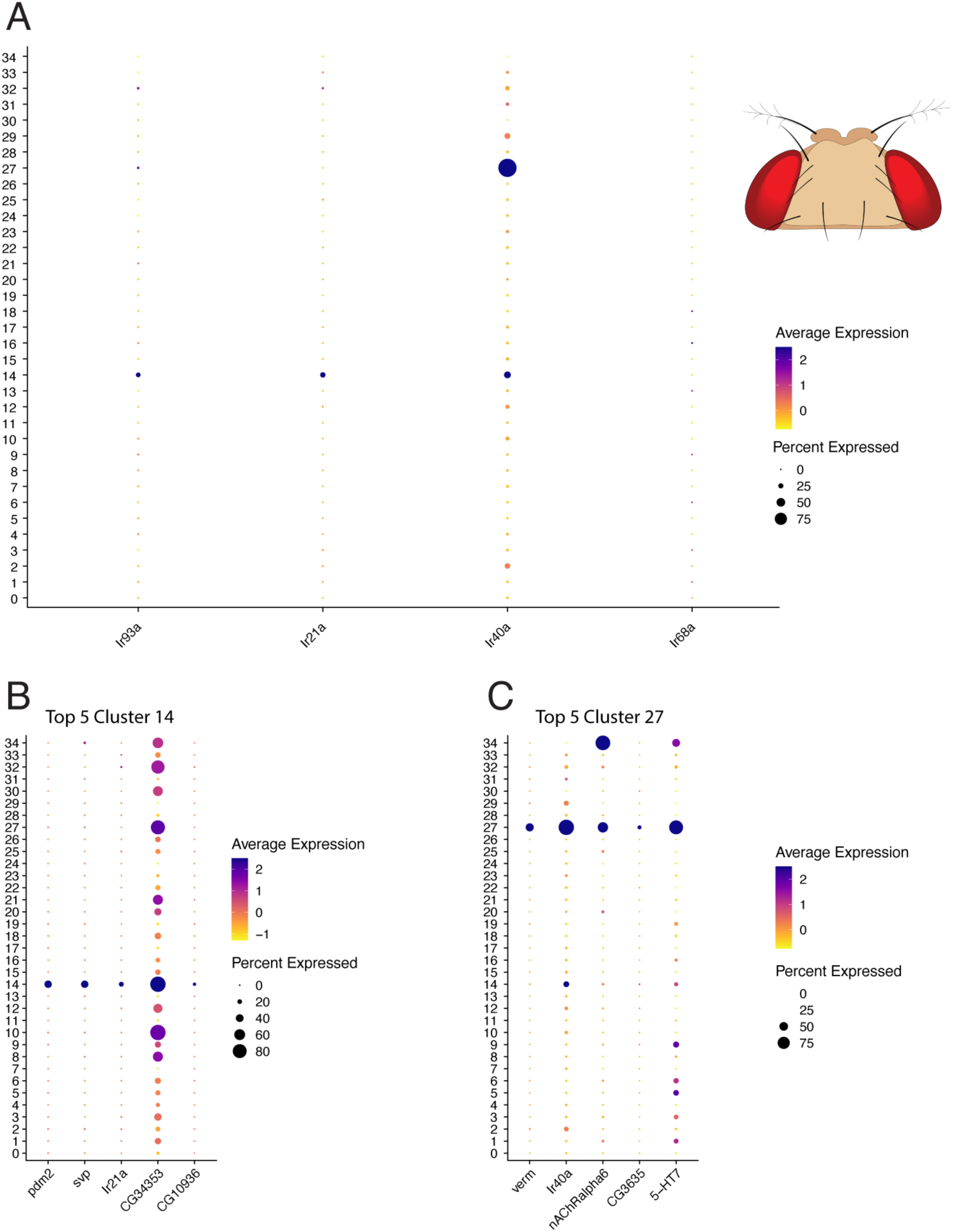
Identification of Hygro- and Thermosensory Neurons in D. melanogaster Antenna (A) Dotplot showing antennal sensory neuron clusters in D. melanogaster, with three clusters expressing hygro- and thermoreceptors (Ir93a, Ir40a, Ir21a, and Gr28b) identified as clusters 14, 27, and 32. (B-D) Dotplot showing expression of top marker genes in clusters 14, 27, and 32.

**Figure 2.**
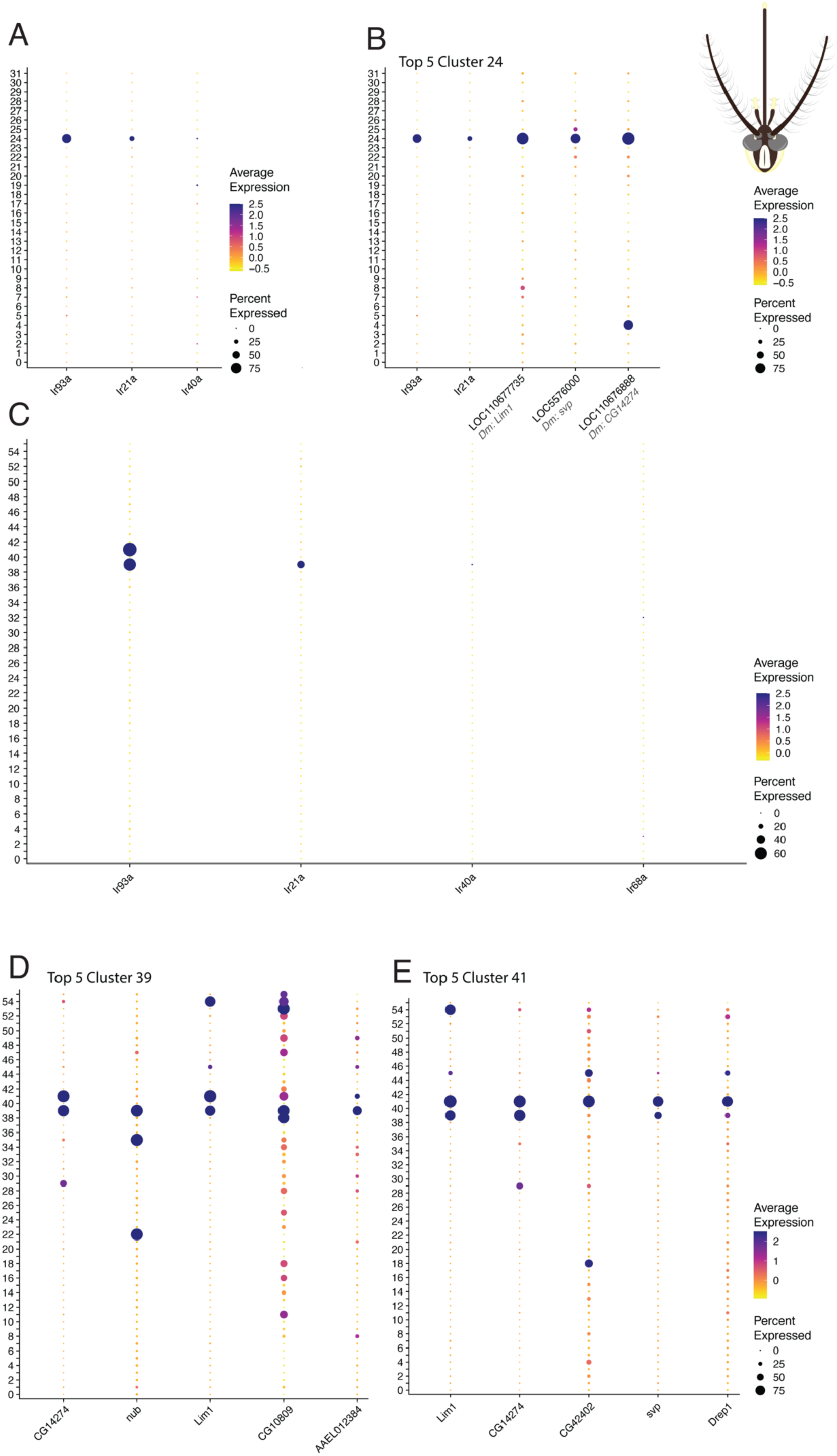
Identification of hygro- and thermosensory neurons in Ae. aegypti antennae. A) Expression levels of hygro- and thermoreceptor genes in Ae. aegypti antennal neurons (dataset from Herre et al). B) Top diQerentially expressed genes in cluster 24. Gray italic text shows D. melanogaster orthologs (dataset from Herre et al). C) Expression levels of hygro- and thermoreceptor genes in Ae. aegypti antennal neurons (dataset from Adavi et al). D) Top diQerentially expressed genes in cluster 39 (dataset from Adavi et al). E) Top diQerentially expressed genes in cluster 39 (dataset from Adavi et al).

In the Herre et al *Ae. Aegypti* dataset, clusters 3 and 45 were excluded from the subsequent analysis due to their ambiguous neuronal marker expression profiles (Supp Figure 2A & B). While these clusters exhibited minor expression of Ir25a, they lacked significant expression of key hygro- and thermoreceptors, including Ir93a, Ir40a, and Ir21a (Supp Figure 3). This absence of relevant receptor expression, combined with their unclear neuronal identity, justified their exclusion from further analyses.

**Figure 3.**
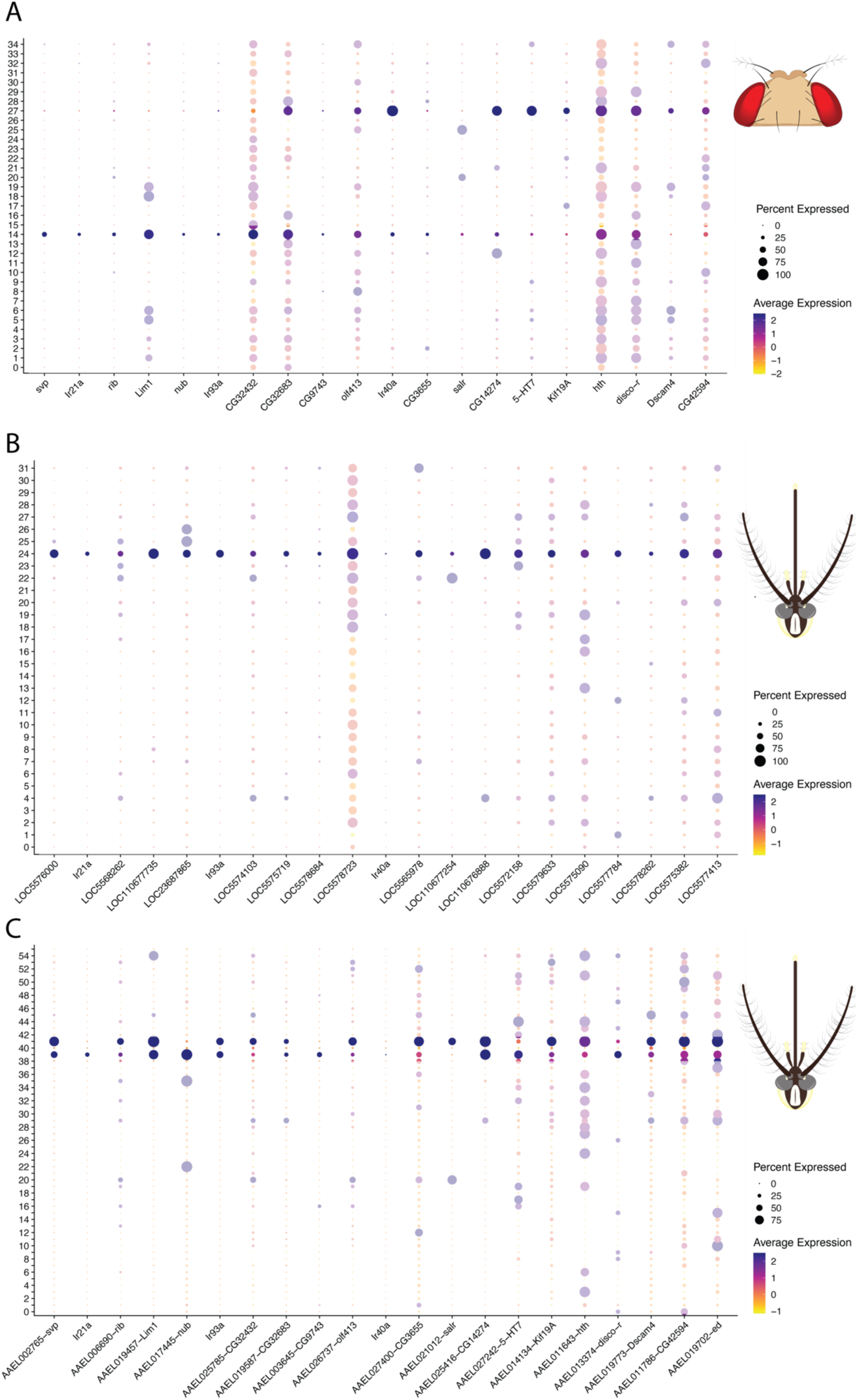
Cross-Species HRN gene comparison in D. melanogaster and Ae. aegypti. A) Expression of shared genes in the HRN/TRN clusters in D. melanogaster. B) Expression of shared genes in the HRN clusters in Ae. aegypti. In grey and italic the D. melanogaster (Dm) orthologue of the Ae. aegypti genes are noted for better comparability (based on cluster 24 from Herre et al dataset). C) Expression of shared genes in the HRN clusters in Ae. aegypti (based on cluster 39 & 41 from Adavi et al dataset).

##### Reprocessing of neuronal clusters and dimensionality reduction

The neuronal clusters were extracted and reprocessed via normalization, PCA analysis and Nearest-Neighbor Clustering. The *D. melanogaster* dataset was re-clustered using 40 principal components (PCs) with a resolution of 0.5. The Herre et al *Ae. aegypti* dataset was analyzed using 40 PCs with a resolution of 1.0. Dimensionality reduction was done by using Uniform Manifold Approximation and Projection (UMAP).

##### Identification of clusters of interest D. melanogaster

To identify which clusters contain the hygrosensory neurons we used expression of the previously described receptors Ir25a, Ir93a, Ir40a, Ir68a, Ir21a and Gr28b.d. Expression of Ir68a was not significant, in line with previous studies^23^.

We then extracted the top5 and top50 marker genes using Seurat *FindMarkers* for the resulting clusters of interest. By cross-referencing those gene lists with our previous transcriptomic study on the HRNs and arista TRNs we assigned cluster identities.

##### Identification of clusters of interest Ae. aegytpi

To identify which clusters contain the hygrosensory neurons we used expression of the preciviously described receptors Ir93a, Ir21a, Ir40a, and Ir68a. No significant expression of Ir68a could be detected. We then extracted the top5 and top50 marker genes using Seurat *FindMarkers* for the resulting clusters of interest.

##### Identification of overlapping genes

To identify genes common to hygrosensory receptor neurons (HRNs) in both species, we compared the top 50 marker genes from the indetified clusters of interest in *D. melanogaster* (clusters 14, 27) with those in *Ae. aegypti* (cluster 24/cluster 39, 41).

To ensure a comprehensive and comparative analysis, we used OrthoDB to identify orthologous genes in both *D. melanogaster* and *Ae. aegypti* for the top 50 marker genes (MG) of each cluster. The resulting lists were then compated as follows:

1. Top 50 MG (cluster 14, 27) from DM x Top 50 MG (cluster 24/cluster 39, 41) from AA as DM orthologues
2. Top 50 MG (cluster 24/cluster 39, 41) from AA x Top 50 MG (cluster 14, 27) from DM as AA orthologues

This resulted in a list of genes that are present in the Top 50 marker genes in both the *D. melanogaster* and the *Ae. aegypti* hygrosensory clusters.

#### Behavioral analysis D. *melanogaster*

##### Animal husbandry and handling

*D. melanogaster* strains were maintained at 25°C on standard *D. melanogaster* food. Humidity levels inside the vials ranged from 70% RH to 90% RH depending on distance to the food source. Prior to the experiment individual *D. melanogaster* were anesthetized on ice and a metal pin was fixed on their thorax using UV curing glue (Helio bond). Subsequently they were starved and desiccated at 5% RH and 21-22 °C for 4 h.

##### Fly strains

*D. melanogaster* strains were obtained from Bloomington Stock Center with the following IDs: #5905 (*w*^1118^), #84446 (w[*]; TI{RFP[3xP3.cUa]=TI}5-HT7[attP]), #358 (w[1118]; nub[2]), #56735 (y[1] w[*]; Mi{y[+mDint2]=MIC}Kif19A [MI12222]), #17177 (w[1118]; P{w[+mC]=EP}Ir21a[EP526]).

##### Behavioral data acquisition

The behavioral data was recorded using the previously described closed-loop dynamic humidity arena^20^. Tethered and desiccated *D. melanogaster* is positioned on a 9 mm plastic ball. The movement of the ball in response to the walking pattern was recorded at 150 fps (Basler acA-1920) and the real-time trajectory is reconstructed using FicTrac^28^. The flies were presented with a step humidity stimulus: 10%RH – 80% RH – 10% RH which each phase being maintained for 500 seconds.

During the experiment humidity and temperature were constantly observed and errors in the delivery of the stimulus or tracking failure led to abortion of the trial. Aborted trials were subsequently removed from the analysis.

##### Data analysis

The obtained real-time trajectories of individuals in response to the humidity stimulus along with humidity level, temperature and time were extracted and used for subsequent analysis.

For noise removal an exponential weighted moving average filter with a span of 50 was applied. Speed data was obtained using the cumulative Euclidean distance between data points and time interval values. For comparability, speed data was normalized and mean speed calculated for 10% RH and 80% RH, respectively (for a more detailed description see^20^).

##### Statistics

As the data was non-normally distributed, statistical significance was determined using a non-parametric bootstrap test with 10,000 resampling iterations. For each comparison, the observed difference in mean speed between humidity levels was calculated. The speed values across the humidity stimuli were pooled and resampled with replacement to generate new datasets. For each resample, the difference in means was computed to form a distribution of bootstrap differences.

The p-value was determined by calculating the proportion of bootstrap differences greater than or equal to the observed difference in absolute value. Comparisons were performed between the three RH setpoints, initial 10% RH, 80% RH and final 10%RH.

## Results

### Identification of HRN populations in fly and mosquito antennal transcriptomes

We used three recently published single-nucleus transcriptomic datasets of antennal neurons: one from *D. melanogaster*^29^ and two independent datasets from *Ae. aegypti*^25,26^. HRN populations were identified in both species’ antennal datasets by examining the expression of genes previously associated with hygrosensation. We therefore searched for presence of Ir93a, Ir40a, Ir68a, Ir21a in both species.

In the *D. melanogaster* antennal transcriptome, we identified three clusters (14, 27, and 32) expressing Ir93a, Ir40a and Ir21a, negative for markers of olfactory neurons (Fig 1A, Supp Figure 4 A). Consistent with previous studies, Ir68a expression was negligible and non-specific throughout the antenna^23^. Cross-referencing cluster-specific markers with our earlier transcriptomic analysis of HRNs revealed that Cluster 14 comprise hygrocool, moist, and arista temperature cells, based on marker gene expression profiles (Figure 1 B, Supp Figure 4 A), while cluster 27 appears to contain dry cells (Figure 1 C, Supp Figure 4 B)^23^.

**Figure 4.**
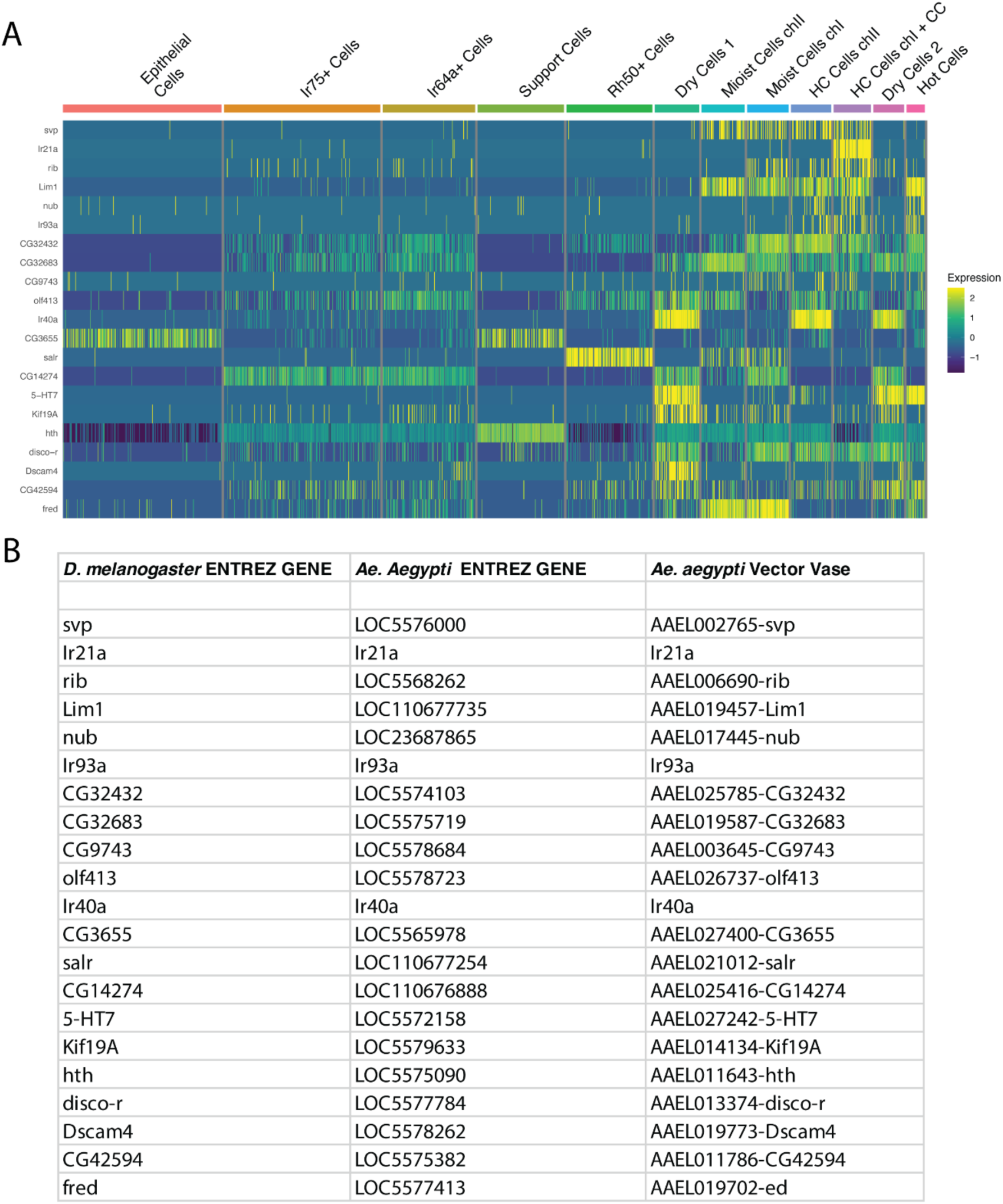
Additional Information for Common Gene List. A) Common genes derived from cross-reference analysis of D. melanogaster and Ae. aegypti Top 5 lists shown in Sacculus and Support Cell sub-clustering from^23^. B) Table displaying the gene IDs of common genes.

Analysis of the two independent *Ae. aegypti* antennal transcriptomes revealed expression of Ir93a, Ir21a, and Ir40a, with Ir40a showing notably low expression levels compared to previous studies (Fig 2). Consistent with our *D. melanogaster* findings, Ir68a expression was insignificant across both datasets. We identified cluster 24 in the Herre et al dataset (Fig 2 A,B) and clusters 39 and 41 in the Adavi et al dataset (Fig 2 C-E) as putative hygrosensory populations based on their expression of these receptors. While the low expression of Ir40a and absence of Ir68a prevented precise classification of specific HRN subtypes, the consistent expression of Ir93a and Ir21a across both datasets, along with their established roles in insect hygrosensation, suggest these clusters represent HRNs in the *Ae. aegypti* antenna.

### Conserved genetic profiles of hygrosensory neurons in dipterans

To determine the conserved genetic profile of HRNs, we compared gene expression profiles between these sensory neurons in *D. melanogaster* and *Ae. aegypti*. By cross-referencing the top 50 marker genes from the HRN clusters in both species (D. melanogaster clusters 14, 27; Ae. aegypti cluster 24 from Herre, and clusters 39, 41 from Adavi), we identified 21 genes, 18 novel and three previously described ionotropic receptors, that likely represent fundamental components of dipteran humidity sensing (Figure 3A – C, Supp Figure 5-7).

**Figure 5.**
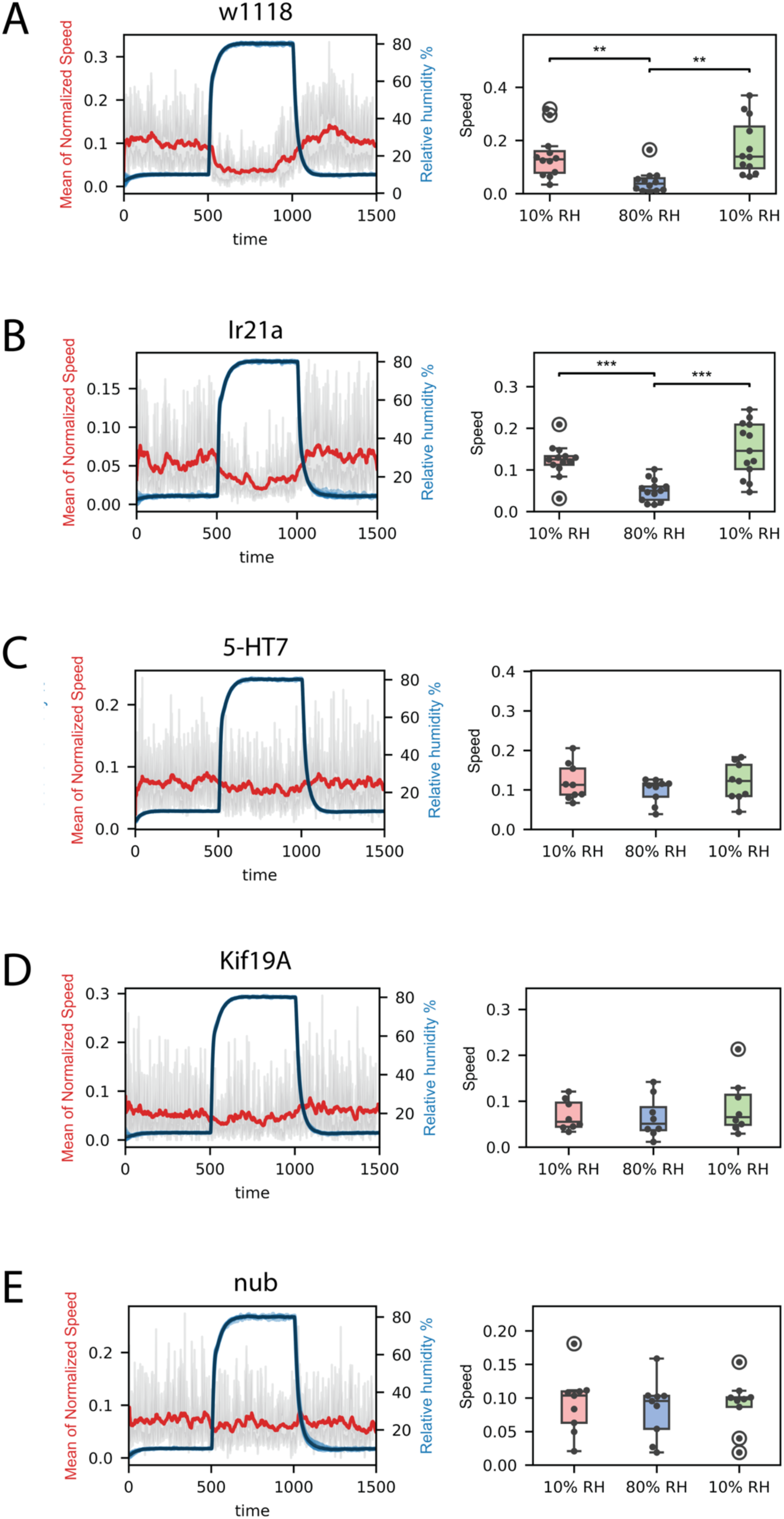
Genetic dissection of humidity-sensing behavior (A-E) Behavioral responses to humidity changes in control and mutant flies. Red lines show mean of normalized speed; gray shading indicates 95% confidence interval. Blue lines represent relative humidity changes (10% → 80% → 10% RH). n = 8-13 flies per genotype. Bootstrapping was used to assess statistical significance. *p < 0.05, **p < 0.01, ***p < 0.001. Box plots show median (line), interquartile range (box), upper and lower quartile (whiskers), individual fly responses (dots) and outliers (circled dots).

These conserved genes fall into distinct functional categories. The largest group consists of transcriptional regulators: the steroid receptor seven-up (svp), the BTB/POZ domain nuclear factor ribbon (rib), LIM homeobox 1 (Lim1), the POU/homeodomain transcription factor nubbin (nub), the zinc finger transcriptional repressor spalt-related (salr), the homeodomain transcription factor homothorax (hth), and the C2H2 zinc-finger transcription factor disco-related (disco-r). The presence of multiple conserved transcription factors suggests a complex, hierarchical regulation of sensory neuron identity in the sacculus and arista.

A second category includes proteins involved in cellular signaling and ion transport: the serotonin receptor 5-HT7, the potassium channel CG42594, the arrestin CG32683, and the already known ionotropic receptors Ir21a, Ir40a and Ir93a. We also identified several enzymes: the dopamine beta-monooxygenase olf413, involved in catecholamine synthesis; CG9743, which shows stearoyl-CoA desaturase activity; and CG3655, which functions as a glycosyltransferase.

Finally, we identified structural and adhesion components include Kinesin family member 19A (Kif19A), a microtubule-associated motor protein, the GPI-anchored proteins CG14274/witty and CG32432, Down syndrome cell adhesion molecule 4 (Dscam4), and friend of echinoid (fred) suggesting the importance of specific cellular recognition and organization in these sensory systems.

For a more detailed understanding of the relationship between specific genes and neuronal subtypes, we conducted an expression analysis using the sub-clustered dataset of sacculus neurons and associated support cells, building upon our previous study’s methodology. This approach allowed for a more nuanced examination of gene expression patterns within these specialized structures^23^. This analysis revealed distinct expression patterns across various sensory neuron subtypes (Fig 4 A, B). In hygrocool and moist cells, we detected expression of the steroid receptor svp. The transcription factor rib exhibited differential expression across hygrosensory neuron subtypes, with notably high levels in hygrocool cells of chamber I and lower expression levels in moist cells. We further observed distinct expression of Lim1 in hygrocool, moist, arista cold, and hot cells, while noting its absence in dry cells. Within temperature-sensitive cells, including hygrocool cells in chambers I and II, as well as arista cold and hot cells, we found nub expression overlapping with Ir93a. Analysis of moist cells showed predominant expression of salr and fred. We detected the serotonin receptor 5-HT7 in dry and arista hot cells, while expression of Kif19A was specific to dry cells. The remaining genes displayed broader expression patterns across sacculus cells, suggesting they may function as more general markers for this neuronal population.

The identification of these 18 novel conserved genes between *D. melanogaster* and *Ae. aegypti* sensory neurons provides insight into the core molecular components required for hygrosensation. The diverse functional categories represented -from transcriptional regulators and signaling molecules to enzymes and structural proteins -suggests that these sensory abilities require the coordinated action of multiple cellular pathways. While some of these genes were previously known to function in sensory systems, others represent novel candidates for future investigation. Their conservation across species, combined with their precise expression patterns in *D. melanogaster* sacculus neurons, indicates they likely play fundamental roles in how insects detect and process environmental humidity information.

### Genetic analysis reveals distinct roles for 5-HT7, nub and Kif19a in hygrosensory system function

To elucidate the functional significance of our identified conserved molecular toolkit in hygrosensation, we performed a humidity behaviour analysis on select genes in *D. melanogaster*. We prioritized genes exhibiting highly specific expression in HRNs across both species and with available viable mutants: Ir21a (hygrocool cells chamber I), 5-HT7 (dry cells chamber I & II), nub (hygrocool cells chamber I & II), and Kif19A (dry cells chamber I & II). This approach allowed us to directly assess the impact of these conserved genes on humidity-sensing behavior. We employed a recently developed dynamic humidity arena^20^ to measure real-time behavioral responses to controlled humidity changes. This fly-on-a-ball system enables precise manipulation of environmental moisture while continuously tracking animal behavior, offering a direct readout of humidity-sensing capabilities. Dehydrated *D. melanogaster* was exposed to humidity transitions ranging from 10% to 80% relative humidity (RH), with their behavioral responses monitored throughout the experiment (Fig 5).

Despite its conservation and expression in hygrocool neurons, Ir21a mutants exhibited typical humidity responses, characterized by increased locomotor activity at low humidity levels and decreased activity at high humidity levels. This pattern mirrors the behavior observed in the control group, indicating potential functional redundancy or compensatory mechanisms in humidity detection (Fig 5 A-B). Specifically, it suggests that hygrocool neurons of sacculus chamber I might not be critical for humidity-guided behaviors under our testing conditions, or that other pathways can compensate for their function. Similar observations have been made for Ir21a mutants in previous reports assessing humidity-guided behaviors^15,17^.

In contrast, loss of the serotonin receptor 5-HT7 and the kinesin Kif19A resulted in significantly impaired humidity-sensing behavior, with individuals showing no response to change in humidity (Fig 5 C, D). While complete loss-of-function mutations in the nubbin (nub) gene are lethal, hypomorphic mutants, which retain partial gene function, are viable. One such mutant, nub^2^, significantly impair humidity-sensing behavior in our behavior assay (Fig 5 E).

Our behavioral analysis reveals differential roles for these conserved genes in humidity sensing, with 5-HT7 and nub demonstrating critical functions in humidity-guided behavior.

## Discussion

### Conserved genes in hygrosensation

Hygrosensation remains one of the few sensory modalities for which the molecular transduction mechanism is not fully understood. Through our comparative analysis, we identified a conserved set of 21 genes associated with hygrosensory neuron function in *D. melanogaster* and *Ae. aegypti*. The conservation of these genes across dipteran species separated by over 200 million years of evolution suggests they represent fundamental components required for humidity sensing. These genes span multiple cellular functions, such as transcription factors, enzymes and membrane-proteins, suggesting humidity detection requires coordinated action across diverse molecular pathways.

Seven conserved transcription factors (svp, rib, Lim1, nub, salr, hth and disco-r) were identified and likely form the regulatory foundation that establishes and maintains hygrosensory neuron identity. Many of these factors have developmental roles, like hth essential for proper antenna formation and segmental identity or Lim1 responsible for specify appendage development^37,38^. However, their continued expression in adult neurons suggests additional roles in maintaining sensory cell identity and function. The transcription factors show distinct expression patterns that align with specific sensory neuron subtypes: salr marks moist cells, while nub is expressed in temperature-sensitive neurons including hygrocool cells and thermal sensors in both chamber I and II. The combinatorial organization of these transcription factors likely creates a regulatory code that defines the molecular and functional properties of each hygrosensory neuron type. The conservation of this transcriptional regulatory network between flies and mosquitoes suggests it constitutes a fundamental molecular framework governing hygrosensory neuron identity and differentiation.

The cell adhesion molecules Fred and Dscam4 show distinct expression patterns in hygrosensory neurons, suggesting specialized roles in the structural organization among these cell types. Fred, which coordinates cellular adhesion and signaling during development^39,40^, is specifically expressed in moist cells where it likely maintains their specialized architecture within the sacculus. Dscam4 is expressed in one subtype of dry cells, likely located to one chamber of the sacculus. Interestingly, previous work has shown that a different family member, Dscam2, marks dry cells in the other chamber, suggesting that different Dscam proteins help establish the distinct organization of dry cell subtypes^23^. Given that Dscam proteins mediate precise neuronal targeting through homophilic adhesion^41^, this complementary expression pattern may be important for maintaining proper circuit architecture in the adult sacculus.

Two conserved GPI-anchored proteins, CG14274/witty and CG32432, potentially play important roles in organizing and regulating humidity receptor complexes. These proteins have well-established functions in modulating ionotropic glutamate receptors (iGluRs) in other contexts: CG14274 mediates receptor aggregation at synapses to ensure proper synaptic transmission^42^, while the C. elegans ortholog of CG32432, called sol-1, acts as a transmembrane AMPAR regulatory protein (TARP) that stabilizes receptor conformations and influences gating properties^43,44^. Given that hygrosensory ionotropic receptors (IRs) share structural similarity with iGluRs, these proteins may serve analogous functions in humidity detection. CG14274/witty could organize IR complexes in hygrosensory dendrites, while CG32432 might modulate their gating properties and stability. The conservation and specific expression of these proteins in hygrosensory neurons suggests they represent fundamental components of the molecular machinery underlying humidity sensing.

The identification of these 18 novel conserved genes between *D. melanogaster* and *Ae. aegypti* sensory neurons provides insight into the core molecular components required for hygrosensation. While some of these genes were previously known to function in sensory systems, others represent novel candidates for future investigation. Their conservation across species, combined with their precise expression patterns in *D. melanogaster* sacculus neurons, indicates they likely play fundamental roles in how insects detect and process environmental humidity information

### Novel genes in hygrosensation

Our behavioral analysis revealed three genes with previously uncharacterized roles in humidity sensing: the serotonin receptor 5-HT7, the kinesin motor protein Kif19A, and the transcription factor nubbin, each providing distinct insights into the cellular mechanisms underlying hygrosensation. 5-HT7 has previously been shown to coordinate sensory input with physiological responses in *D. melanogaster*, specifically in translating gustatory detection into changes in digestive function by activating enteric neurons to regulate crop contractions^45^. This fits with broader evidence that serotonergic systems can transform acute sensory detection into longer-term physiological changes, allowing animals to make anticipatory responses to environmental conditions^46^. Given these findings, and serotonin’s established role as a diuretic hormone in several insect species^47^, 5-HT through 5-HT7 may play a similar integrative role in humidity sensing, potentially modulating the excitability of hygrosensory neurons to fine-tune their responses to environmental moisture changes. As the highest-affinity serotonin receptor in *D. melanogaster*, 5-HT7 is particularly well-suited to detect subtle changes in circulating serotonin levels, suggesting it could integrate both local and systemic serotonergic signals to modulate hygrosensory responses^48^. The dual role of 5-HT7 in both regulating feeding and humidity tracking also raises the possibility that these sensory modalities are coordinated to help flies maintain water balance, particularly important given that feeding state and humidity levels both impact desiccation risk. Further experiments will be needed to determine how 5-HT7 acts directly in the dry cells and how it modulates their activity.

The significant behavioral deficits observed in Kif19A mutants suggest a critical role for this kinesin in hygrosensation, potentially through its involvement in intracellular transport or maintenance of cilia structure, illustrating the importance of cellular infrastructure in sensory neuron function. Kif19A, a plus-end directed kinesin-8 family motor protein, exhibits microtubule-depolymerizing activity and localizes specifically to ciliary tips^49^. Studies in mice have demonstrated that loss of Kif19A leads to abnormally elongated cilia that cannot function properly^49^. As ciliary length and architecture is a critical determinant of sensory function, Kif19A ’s control of ciliary length in dry cells may be essential for proper detection of humidity changes^50,51^. Furthermore, ciliary membrane composition and receptor localization are tightly regulated processes essential for sensory function^52^. As a plus-end directed motor, Kif19A may also play a role in trafficking sensory components, like Ionotropic receptors, to ciliary tips. However, the exact relationship between microtubule structure and hygrosensory function requires further investigation to fully understand the role of Kif19A in this sensory system.

The hypomorphic nub2 mutant, which retains partial gene function of nubbin and therefore viability, exhibits significantly impaired humidity-sensing behavior in our assay. As a POU-domain transcription factor, nub likely coordinates multiple aspects of sensory neuron development and maintenance. Recent work has shown that nub functions redundantly with its paralog pdm2 in wing development, where both genes respond to a shared enhancer element called R11F02 to maintain wing identity^53^. This enhancer is also specifically active in HRNs and TRNs in the adult antenna^50^ suggesting a similar role of nub in sensory neurons, potentially maintaining the expression of essential sensory components. nub could coordinate multiple aspects of these specialized neurons, from maintaining expression of critical receptors to regulating ion channels necessary for proper sensory transduction.

These expanded roles for 5-HT7, nub and Kif19A highlight how receptors, transcriptional regulation and cellular transport cooperate to establish and maintain humidity sensing. The conservation of these components across species suggests they represent fundamental requirements for hygrosensory neuron function, providing new insights into the cellular machinery underlying environmental sensing in insects.

### Concluding remarks

Our integrative analysis of hygrosensory neurons across two phylogenetically distant dipteran species has revealed a conserved molecular toolkit essential for hygrosensation. By combining comparative transcriptomics with targeted behavioral analysis, we have uncovered multiple layers of cellular regulation in these specialized neurons. Beyond the previously characterized ionotropic receptors, we identified novel components spanning diverse cellular functions: the serotonin receptor 5-HT7 suggesting potential neuromodulatory mechanisms, the transcription factor nubbin indicating requirements for sustained gene regulation, and the motor protein Kif19A pointing to a role for specialized cellular transport. The conservation of these components between species separated by over 200 million years of evolution, together with their specific expression patterns in different sensory neuron subtypes, suggests fundamental organizational principles in HRNs. This approach, combining systematic genomic comparison with targeted functional analysis, provides a framework for investigating sensory system evolution across species.

## Acknowledgements

We wish to thank the NBIS for their support on this project. The authors thank Marcus Stensmyr for comments on discussion of this project.

## Authors contributions

KC and AE designed the study. KC performed the transcriptomic study with input from JR and AC. GG performed and analyzed the behavioral experiments. KC and AE wrote the manuscript with input from all authors.

## Supplementary Figures

**Supplemental Figure 1.**
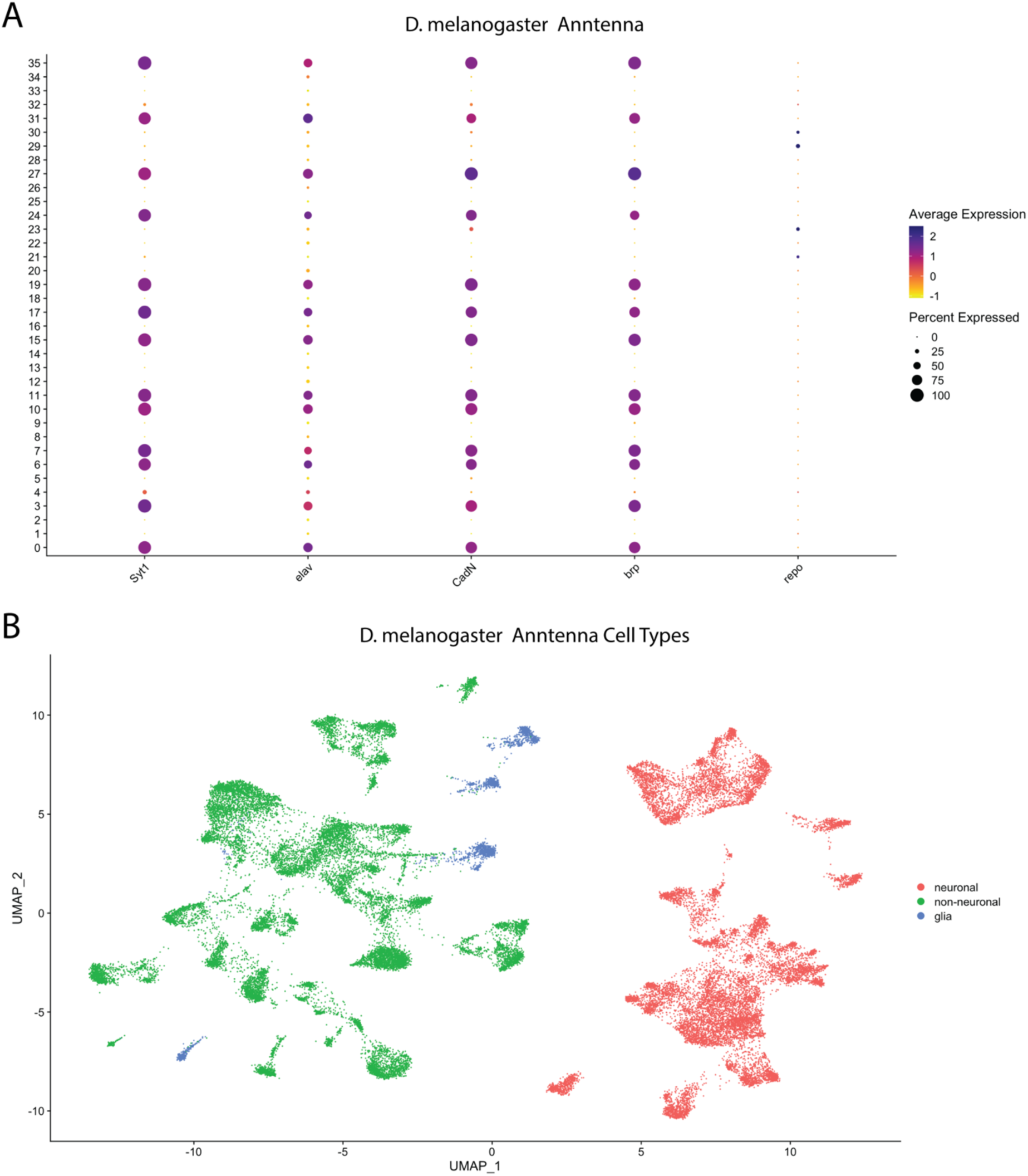
Identification of neuronal and glia clusters in the D. melanogaster antenna. A) Dot plot showing expressions of the neuronal markers Syt1, elav, CanN, brp and the glial marker repo. B) UMAP projection showing the allocation of cluster identities into the cell types neuronal, non-neuronal, glia.

**Supplemental Figure 2.**
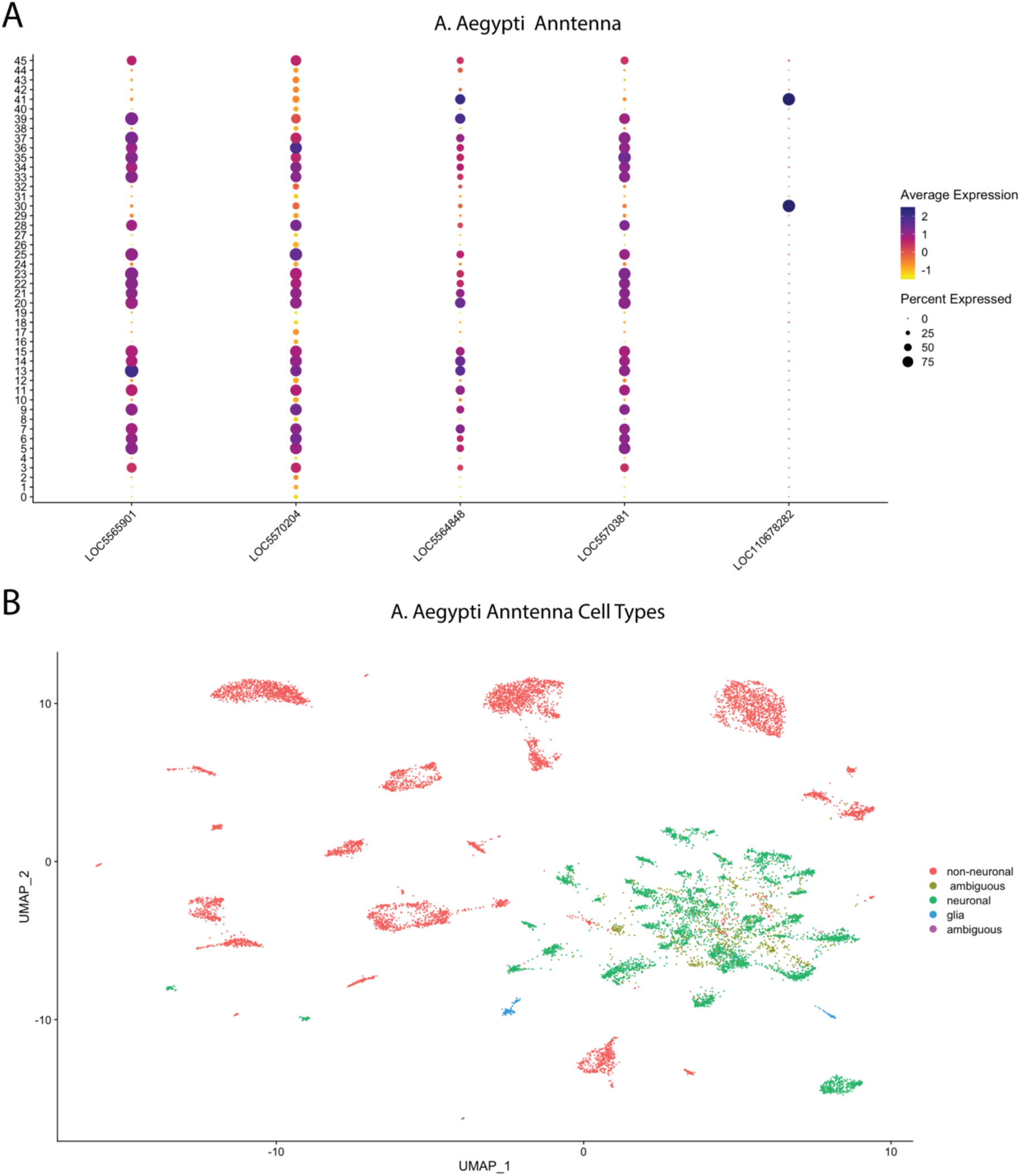
Identification of neuronal and glia clusters in the A. aegytpi antenna. A) Dotplot showing expressions of the neuronal markers LOC5565901 (orthologue to Syt1), LOC5570204 (orthologue to elav), LOC5564848 (orthologue to CadN), LOC5570381 (orthologue to brp), LOC110678282 (orthologue to repo). B) UMAP projection showing the allocation of cluster identities into the cell types neuronal, non-neuronal, glia. Based on marker expression cluster 3 and 45 were classified as “ambiguous”.

**Supplemental Figure 3.**
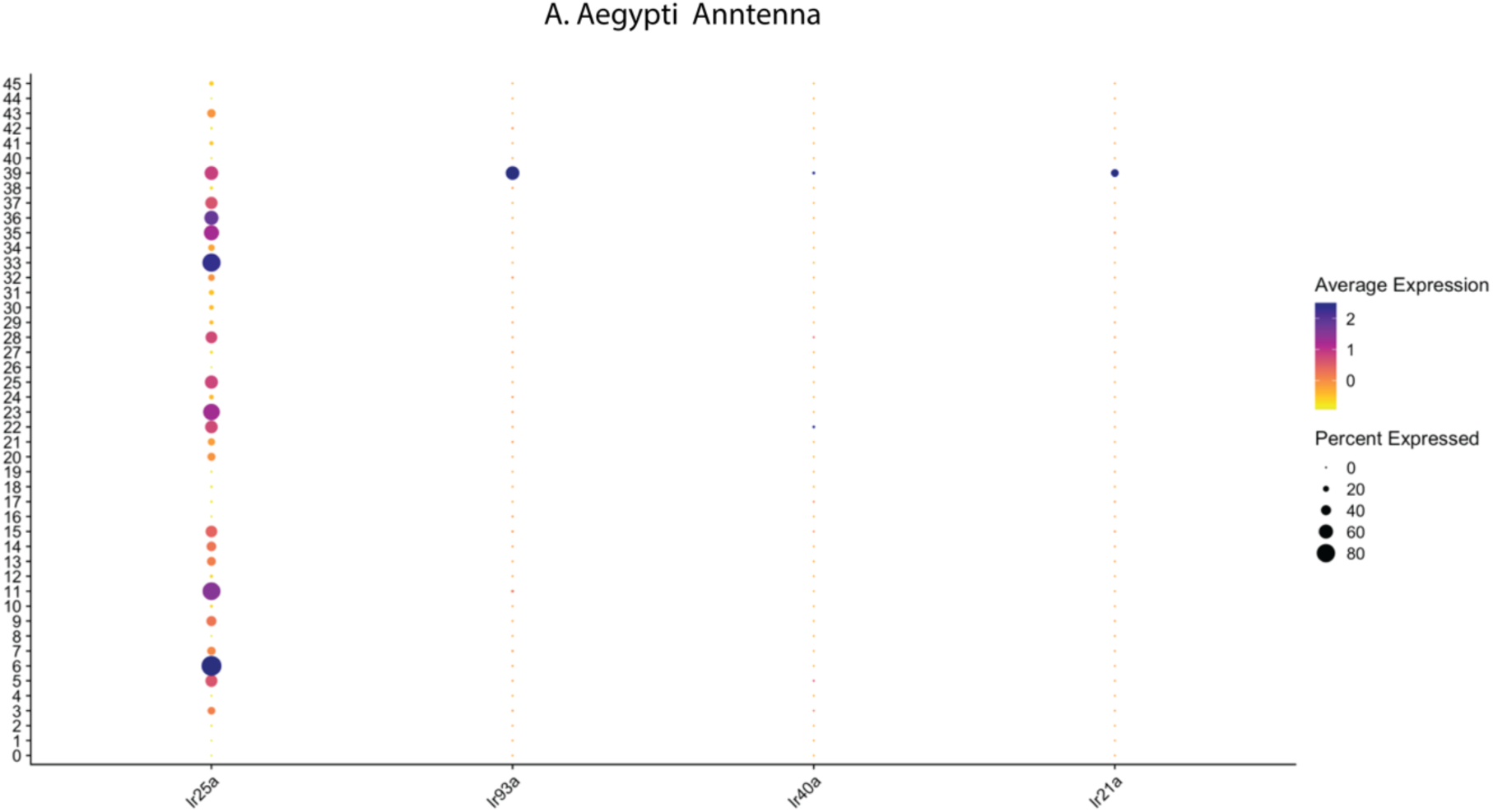
Expression of the hygro- and thermoreceptors Ir25a, Ir93a, Ir40a and Ir21a in the A. aegyti antennal dataset. Absence of relevant expression in clusters 3 and 45 justifies their exclusion from further analysis.

**Supplemental Figure 4.**
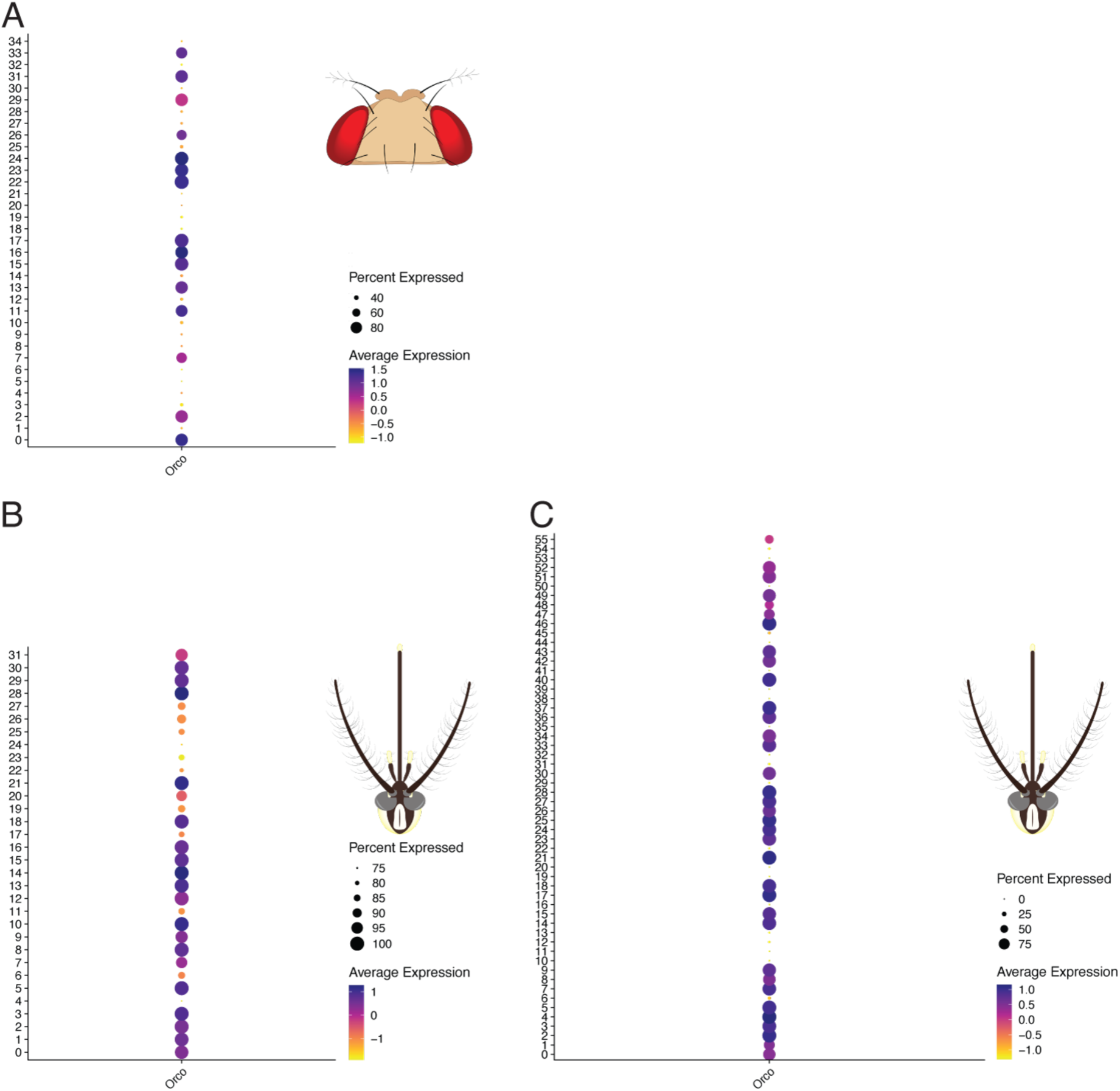
A) Expression of Orco in antennal neurons in D.melanogaster B) Expression of ppk, ppk26 and kn in D. melanogaster antennal neurons. The identified HRN and arista TRN clusters are characterized by an absence of Orco.

**Supplemental Figure 5.**
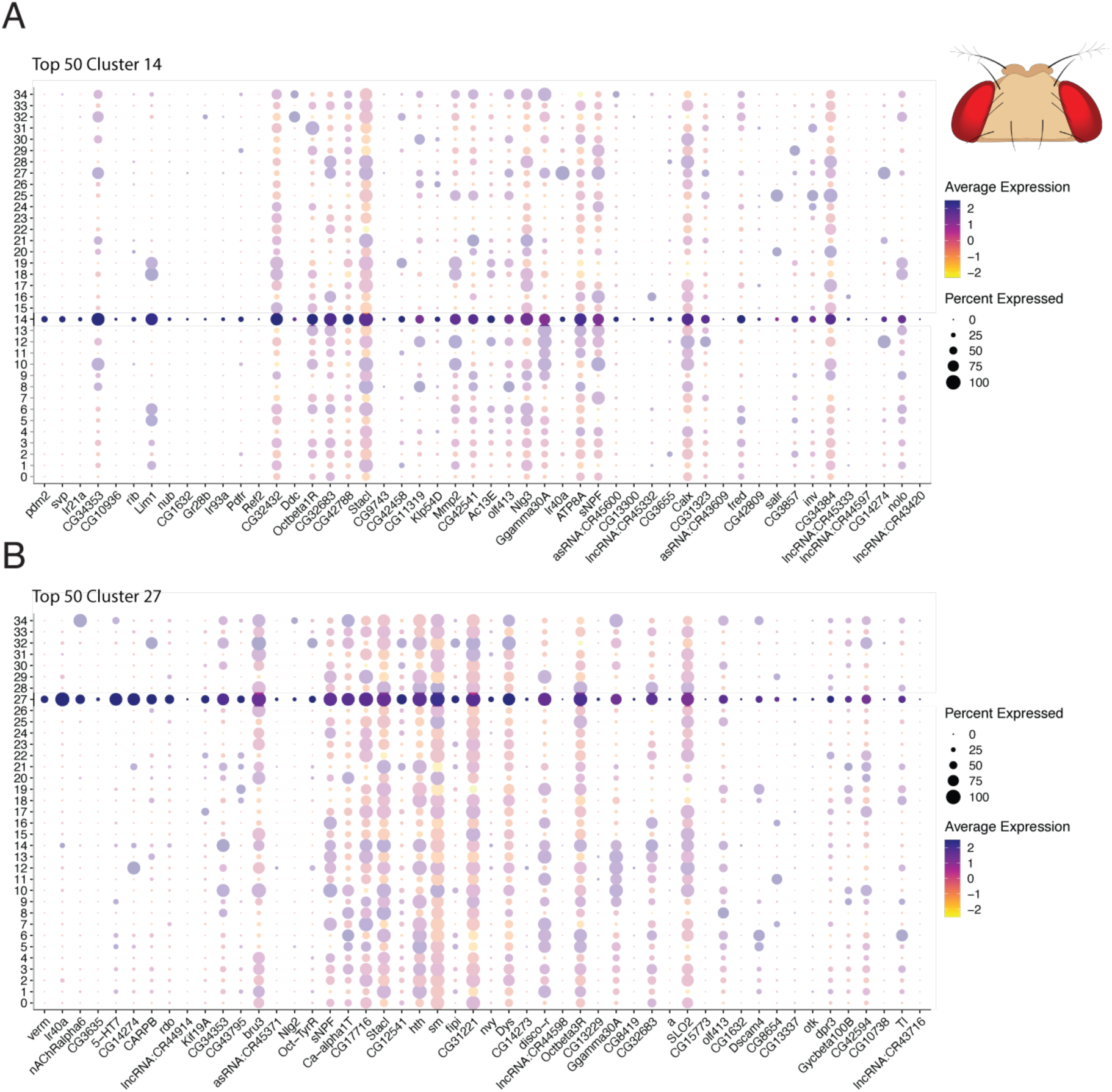
Top 50 markers genes in the antennal neurons of D. melanogaster for A) Cluster 14, B) Cluster 27 and C) Cluster 32.

**Supplemental Figure 6.**
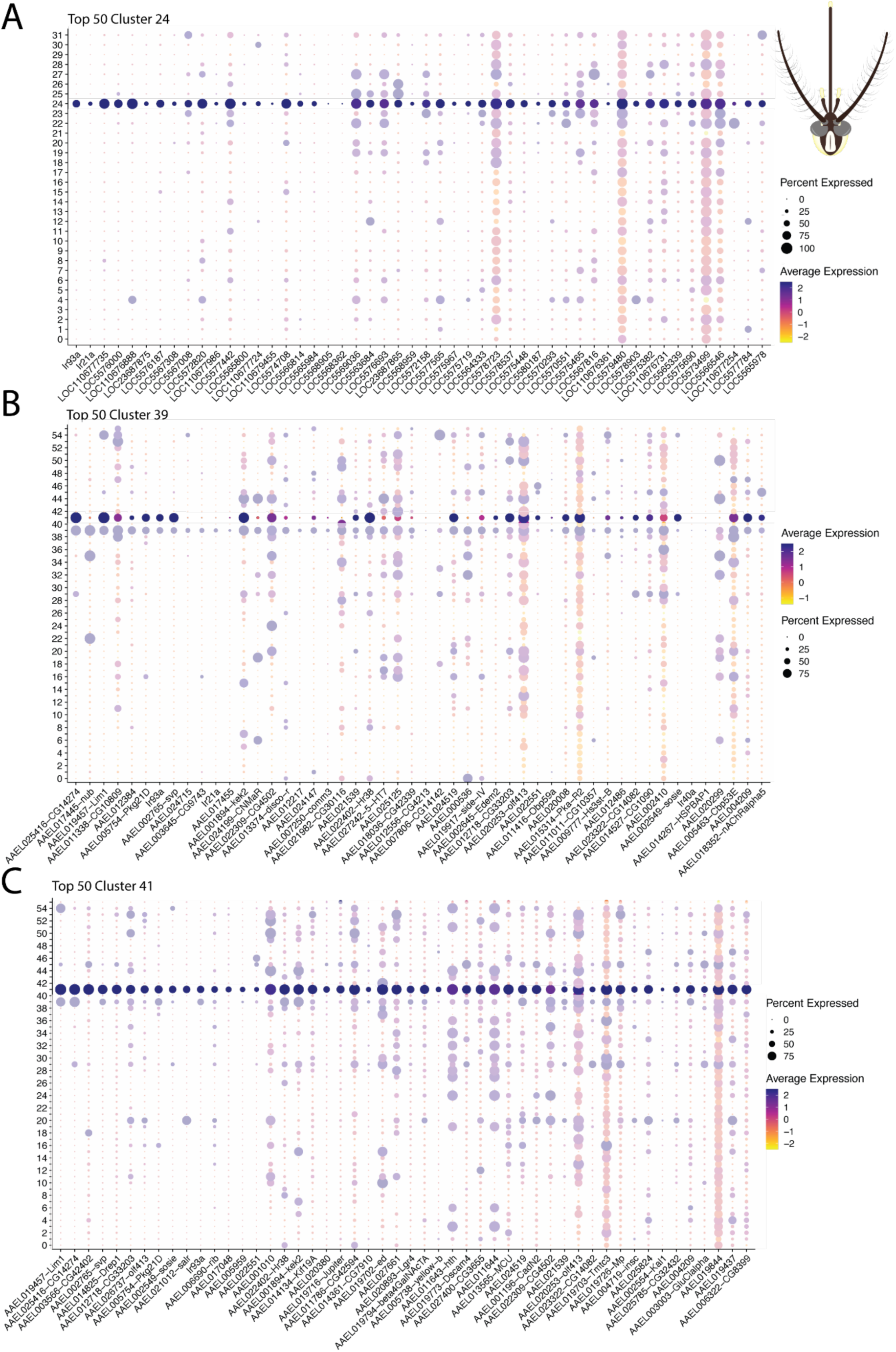
Top 50 markers genes in the antennal neurons of Ae. aegypti for A) Cluster 24 (Herre et al), b) Cluster 39 (Adavi et al, 2024), C) Cluster 41 (Adavi et al, 2024).

